# Emerging Hantaviruses in Central Argentina: first case of Hantavirus Pulmonary Syndrome caused by Alto Paraguay Virus and a novel orthohantavirus in *Scapteromys aquaticus* rodent

**DOI:** 10.1101/2021.06.13.448262

**Authors:** Carla M Bellomo, Daniel Alonso, Tamara Ricardo, Rocío Coelho, Sebastián Kehl, Natalia Periolo, Natalia Casas, Laura Cristina Bergero, María Andrea Previtali, Valeria Paula Martinez

## Abstract

Orthohantaviruses are emerging rodent-borne pathogens that cause Hantavirus Pulmonary Syndrome in humans. They have a wide range of rodent reservoir hosts and are transmitted to humans through aerosolized viral particles generated by the excretions of infected individuals. Since the first description of HPS in Argentina, new hantaviruses have been reported throughout the country, most of which are pathogenic to humans.

We present here the first HPS case infected with Alto Paraguay virus reported in Argentina. Until now, Alto Paraguay virus was considered a non-pathogenic orthohantavirus since it was identified in a rodent, *Hollochilus chacarius*. In addition to this, with the goal of identifying potential hantavirus host species in the province of Santa Fe, we finally describe a novel orthohantavirus found in the native rodent *Scapteromys aquaticus*, which differed from other hantaviruses described in the country so far.

Our findings implicate an epidemiological warning regarding these new orthohantaviruses circulating in Central Argentina as well as new rodent species that must be considered as hosts from now on.

## Introduction

Hantavirus Pulmonary Syndrome (HPS) is an emerging zoonotic disease transmitted to humans through inhalation of aerosolized viral particles generated by the excretions of infected rodents, which act as reservoir hosts in nature [1]. The genus *Orthohantavirus* (family *Hantaviridae*, subfamily Mammantavirinae) includes rodent-borne HPS agents and other viruses not yet associated with human diseases. They are enveloped, single-stranded negative-sense RNA viruses, and their genome is divided in 3 segments: small (S), medium (M), and large (L) [2]. The clinical course of HPS generally progresses through three phases: prodromal, cardiopulmonary and convalescent. Clinical manifestations can vary from a mild hypoxemia to a respiratory failure with cardiogenic shock [3]. HPS was first described in 1993, due to an outbreak in the Four Corners region of the United States [4].

In South America, there are 5 species of orthohantavirus: Andes, Cano Delgadito, Laguna Negra, Maporal and Necocli, including more than 10 named viruses [5]. Andes virus (ANDV) was the first HPS agent described in Argentina [6]. Seven pathogenic ANDV-like viruses were subsequently characterized in 4 endemic regions in the country. To date, in Central Argentina, HPS cases are associated with 3 ANDV-like viruses: Buenos Aires virus, Plata virus, and the officially named Lechiguanas virus [7; 8]. All these viruses are believed to be transmitted to humans by the same reservoir host, *Oligoryzomys flavescens* [9, 10]. In the same region, 2 viruses that are currently not associated with human diseases, Maciel and Pergamino viruses, were characterized in the native rodents *Necromys sp*. and *Akodon azarae*, respectively [11]. In the present work, we report an HPS case infected with Alto Paraguay virus (APV), an orthohantavirus never before reported in the country, and previously considered as non-pathogenic. The case was reported beyond the northernmost HPS endemic area of Central Argentina. Analysis of rodent populations in the periphery of the known HPS endemic area led us to the identification of a novel orthohantavirus in the native rodent *Scapteromys aquaticus*. These findings implicate an expansion of the known HPS endemic area in Central Argentina and a greater diversity of orthohantavirus in the country.

## Methods

### Case definition and epidemiological data

HPS case was defined as a patient who resides or reports a recent travel history to an endemic region, characterized in the initial stage by persistent fever (>48 hours), headache, myalgia, gastrointestinal manifestations (abdominal pain, vomiting and/or diarrhea), and a marked decrease in platelet count [8]. After this short prodromal phase, manifestations of respiratory compromise are common. The clinical history and laboratory data from the patient were retrieved from medical records of healthcare institutions from the province of Santa Fe. The patient was a young male residing in Tostado (−29.2333, -61.7667), a town of 15,500 inhabitants, distributed over 390,800 hectares mainly dedicated to livestock farming and agriculture, in the ecoregion called Dry Chaco [12]. The total number and distribution of HPS cases in the province of Santa Fe from 1996 to 2020 was obtained from our own database and from Sistema Nacional de Vigilancia de la Salud (Ministerio Nacional de Salud y Desarrollo Social). Confirmed cases, those with IgM and IgG, or IgM and RT-PCR, are shown in Fig 1.

**Figure 1.**
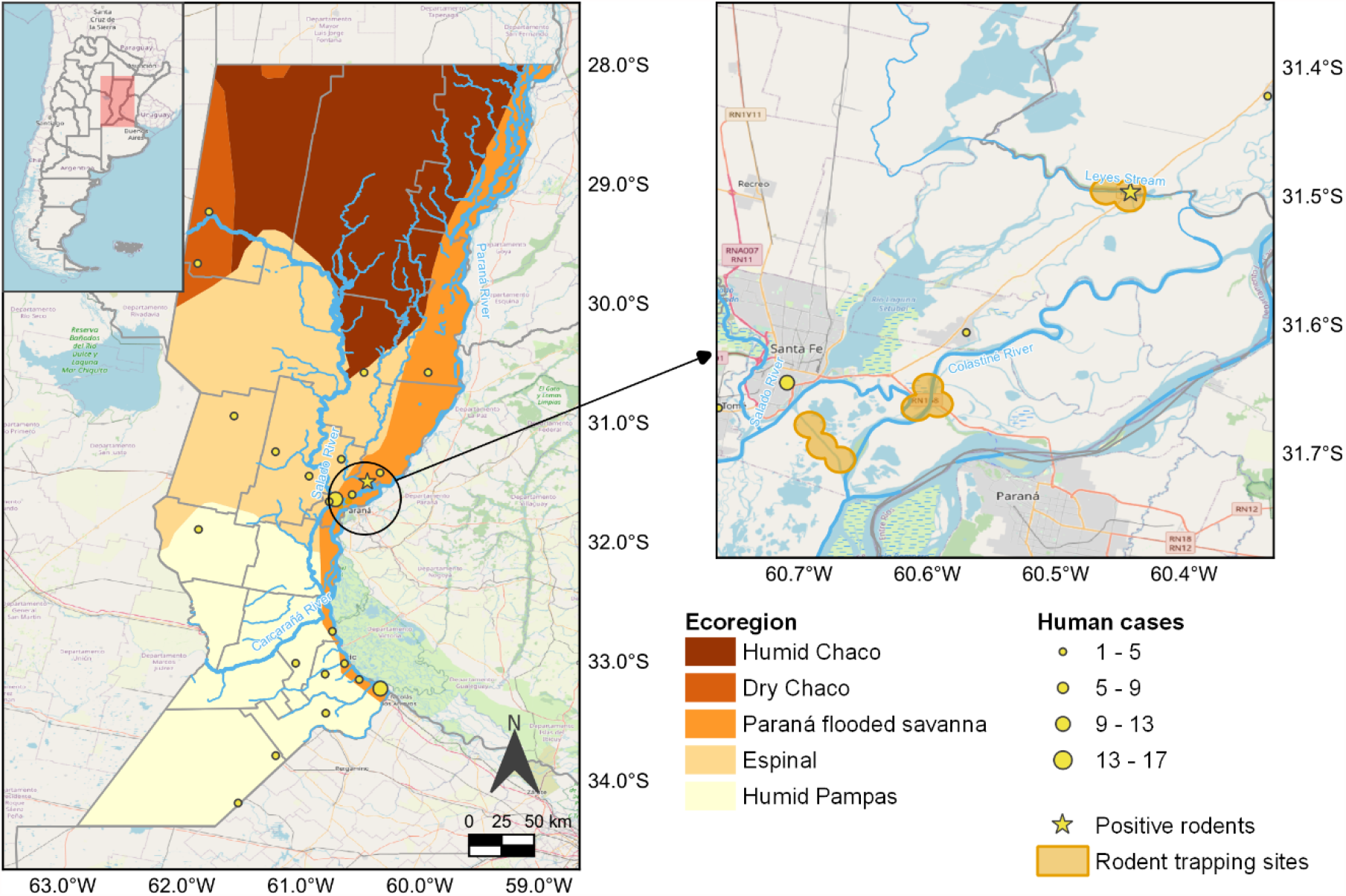
Hantavirus Pulmonary Syndrome cases distribution in the province of Santa Fe, Argentina, from 1996 to 2020. The small map displays a zoomed view of the city of Santa Fe and the 3 riverside communities where the rodent sampling was conducted (Sep. 2014-Oct. 2015). Map created using QGIS 3.14 Pi (QGIS Development Team). Vector layers downloaded from <u>Biodiversity Information System of the National Parks Administration, Argentina (SIB)</u>, <u>Base of Human Settlements of the Argentine Republic (BAHRA)</u> and <u>National Geographic Institute (IGN)</u>. Base map from <u>OpenStreetMap</u>

### Laboratory confirmation

Two days after the onset of symptoms, and due to the ongoing pandemic of Coronavirus (SARS-CoV-2), the patient was subjected to a nasopharyngeal swab (NFS) to determine infection by SARS-CoV-2 by real time reverse transcription-polymerase chain reaction (RT-qPCR) by the National Reference Laboratory for Respiratory Viruses (Instituto Nacional de Enfermedades Infecciosas – ANLIS “Dr. C. G. Malbrán”). The serum sample of the patient was processed in our laboratory (National Reference Laboratory for Hantavirus) to confirm hantavirus infection. We performed enzyme-linked immunosorbent assays (ELISA) to detect ANDV-specific immunoglobulin M (IgM) and immunoglobulin G (IgG) in serum as previously described [13]. Hantavirus genome detection was carried out from serum and NFS by total RNA extraction using Trizol according to the manufacturer’s protocol (Thermo Fisher Scientific), followed by endpoint RT-PCR.

### Rodent trapping and seroprevalence study

With the goal of identifying potential hantavirus host species in the province of Santa Fe, we processed samples from rodents captured during a ecoepidemiological study of leptospirosis that involved two springs (Sept-Oct 2014 and 2015) and one autumn (Mar-Apr 2015) in three communities of the Paraná flooded savanna (Fig 1). Each community was located on the banks of a water course (i.e., Leyes stream, Colastiné river, and the port accessibility channel; Fig. 1). Detailed information on the study protocol can be found in Ricardo *et al*. [14, 15]. Briefly, each community was splitted into 3 study sites based on their level of human disturbance, and 25 trapping stations (consisting of a Sherman trap and a live-cage trap) were set per site during 3 consecutive nights. Blood samples were collected by cardiac puncture and tissue samples by systemic necropsy. Morphometric and reproductive features, body condition score [16] and presumptive species were recorded for each captured animal. Rodent species were subsequently confirmed by inspection of skull morphology or by molecular identification through mitochondrial genes from liver and/or lung samples [15]. Samples were stored in liquid nitrogen during the trapping session and for their transportation to the laboratory. There, they were stored in a -701 freezer and finally moved to a -20⍰ freezer.

Serum samples were tested by IgG ELISA as previously described [17]. Briefly, serum samples were four-fold diluted from 1:400 to 1:102,400 and incubated in microplates coated with recombinant nucleoprotein from ANDV, Seoul virus or a recombinant control protein, followed by peroxidase-conjugate antibodies against mouse or rat IgG antibodies respectively. After the incubation with TMB, the optical density (OD) was measured at 450 nm, and the ΔOD was obtained subtracting the corresponding OD of the same sample incubated with control recombinant antigen. Samples were considered positive if ΔOD values were greater than 0.3. IgG titers were determined by endpoint dilution and calculated as the highest serum dilution in which OD values were still positive. Associations between seroprevalence and rodent species or environmental settings were assessed using Pearson’s Chi-squared test or Fisher’s exact test.

### Viral Genetic characterization and phylogenetic analysis

For viral genetic characterization the RNA extracted from the patient’s serum (see above) and from lung and/or heart tissues of seropositive rodents were used to amplify viral genome by RT-PCR using the One Step RT-PCR kit (QIAGEN) followed by nested or heminested PCRs (Taq Pegasus, PB-L Productos Bio Lógicos and/or Platinum™ Taq DNA Polymerase, Invitrogen). Initial screening was performed by using primers specific for the S segment as described [18]. Additional primers were selected to obtain sequences spanning the entire M-segment coding region.

Amplicons were purified by agarose gel electrophoresis and sequenced directly using the BigDye Terminator v3.1 Cycle Sequencing Kit on an ABI Prism R3500 (Applied Biosystems, Foster City, CA.). Twenty six previously published sequences, that were obtained from public repositories (GenBank), and the samples sequenced in this study, were aligned by ClustalW in MEGA version X [19]. For the phylogenetic analysis, the determination of the evolutionary model was performed by J Model Test 2.1.416. The selected model was General Time Reversible (GTR), with discrete Gamma distribution (+G) and by assuming that certain fractions of sites are evolutionarily invariable (+I). This analysis involved a total of 1,877 and 3,555 positions, for S- and M-segments, respectively in the final dataset. Bayesian phylogenetic reconstruction was realized using the algorithm Metropolis-coupled Markov chain (MCMC) method available in the BEAST v1.8.4 package [19]. We used Tracer v.1.6 to check for convergence (i.e., an estimated sample size >200 for all relevant parameters). The TreeAnnotator program was used to generate a Maximum Clade Credibility (MCC) tree.

## Results

### HPS case detection

In the context of the COVID-19 pandemic, a 15-year-old male patient attended a hospital looking for medical assistance manifesting fever, general weakness, headache, cough and myalgia. As COVID-19 was first suspected, the patient was admitted to the local hospital, in Tostado, Santa Fe province. A serum sample and a NFS were obtained at admission, 2 days after onset of symptoms. The patient showed a mild clinical presentation with no severe respiratory manifestations, additionally, the computed tomography screening exhibited a picture compatible with COVID-19. However, the patient improved fast and was dismissed after a few days. Clinical laboratory findings at admission are shown in Table 1. The values are concordant with those found in patients with mild disease in ANDV-HPS at early stages [20]. The patient reported no history of travel and manifested no previous contact with any confirmed COVID-19 case, and then HPS was considered as differential diagnosis. Although the residence place of the patient was not considered as an endemic zone for HPS, a previous case was reported in the area 5 months before (Sistema Nacional de Vigilancia de la Salud, Ministerio Nacional de Salud y Desarrollo Social).

**Table 1.** Laboratory findings in a Hantavirus Pulmonary Syndrome patient at day 2 after the onset of symptoms.

| Parameter | Value | Reference values | Interpretation |
| --- | --- | --- | --- |
| White Blood Cells | 5800 cells/mm<sup>3</sup> | 4500 – 11500 cells/mm<sup>3</sup> | Normal |
| Neutrophiles | 5046 cells/mm<sup>3</sup> | 1300-6000 cells/mm<sup>3</sup> | Normal |
| Lymphocytes | 580 cells/mm<sup>3</sup> | 1500-3500 cells/mm<sup>3</sup> | Lymphopenia |
| Monocyte | 174 cells/mm<sup>3</sup> | 300-900 cells/mm<sup>3</sup> | Normal |
| Platelets | 83,000 cells/mm<sup>3</sup> | 150,000-450,000/mm<sup>3</sup> | Low |
| Hemoglobin | 15.1g/dl | 14.4-17.4 g/dl | Normal |
| Hematocrit | 46.7% | 40-54% | Normal |
| Creatinine | 0.95 mg/dl | 0,52-1,04 mg/dl | Normal |
| Urea | 17mg/dl | 12-54 mg/dl | Normal |
| Alkaline phosphatase | 32 IU/L | 74 to 390 IU/L | Low |
| AST | 45U/l | 17-59 U/L | Normal |
| ALT | 20U/l | 21-72 U/L | Normal |
| Sodium | 142 mEq/l | 135-145 mEq/L | Normal |
| Potassium | 3.6 mEq/l | 3.5-5.1 mEq/L | Normal |
| Total bilirubin | 6 mg/dl | <1mg/dl | High |

RT-PCR for coronavirus and hantavirus in the NFS sample were both negative. However, as NFS is not adequate for routine hantavirus diagnosis, a serum sample was subjected to ELISA and RNA extraction. The serum sample had detectable levels of IgM and IgG antibodies to hantavirus, with high IgM titers (6,400) and low IgG titers (100). For viral genetic characterization, two partial fragments were successfully amplified from the S and M segments, 320pb and 741pb respectively, corresponding to positions 34-356 and 2,219-2,960 from GenBank accession numbers AF324902 and AF324901, respectively. Pairwise nucleotide sequence comparisons with previously published sequences showed low identities with all the hantavirus previously characterized in Argentina. The highest nucleotide identities, 98.6% for the S-segment and 95.3% for the M-segment (Table 2), were found with APV, an orthohantavirus that was previously described in western Paraguay associated with *Holochilus chacarius*. Amino acidic comparison showed 99 and 100% identities for N and G2 proteins respectively with the same virus.

**Table 2.** Pairwise nucleotide and amino acid identity percentages between virus sequences amplified from Hantavirus Pulmonary Syndrome case, Santa Fe province, Argentina and selected American orthohantaviruses.

| Virus | Country | S-segment |  | M-segment |  |
| --- | --- | --- | --- | --- | --- |
|  |  | Nucleot. | Amin. | Nucleot. | Amin. |
| Jabora virus | Brazil | 77.5 | 74.4 | 79.6 | 96.1 |
| Alto Paraguay virus | Paraguay | **98.6** | **95.5** | **95.3** | **100.0** |
| Anajatuba virus | Brasil | 88.4 | 74.4 | NA | NA |
| **Andes virus** | Argentina | 86.8 | 74.4 | 79.8 | 93.9 |
| Araracuara virus | Brasil | 81.8 | 71.8 | NA | NA |
| Bermejo virus | Argentina | 84.3 | 74.4 | 78.4 | 93.9 |
| Buenos Aires virus | Argentina | 86.1 |  | NA | NA |
| **Caño Delgadito virus** | Venezuela | 77.1 | 76.9 | 76.3 | 92.6 |
| **Castelo Do Sonhos virus** | Brazil | 85.8 | 71.8 | 79.8 | 94.4 |
| **Choclo virus** | Panamá | 86.8 | 74.4 | 79.1 | 90.9 |
| **El Moro Canyon virus** | United States | 80.4 | 69.2 | 77.5 | 84.35 |
| IP16 virus | Paraguay | 79.7 | 74.4 | NA | NA |
| Itapúa virus | Paraguay | 85.0 | 74.4 | NA | NA |
| Juquitiba virus | Arg. / Brazil | 84.7 | 74.4 | 79.8 | 96.1 |
| **Laguna Negra virus** | Argentina/<br>Parag. / Brazil | 88.0 | 71.8 | 81.0 | 94.0 |
| **Lechiguanas virus** | Argentina | 84.6 | 74.4 | 78.7 | 94.8 |
| Maciel virus | Argentina | 81.8 | 71.8 | 77.5 | 92.2 |
| **Maporal virus** | Venezuela | 84.4 | 74.4 | 80.5 | 91.3 |
| **Necoclí virus** | Colombia | 84.0 | 74.4 | 81.6 | 92.6 |
| **Oran virus** | Argentina | 85.7 | 74.4 | 79.2 | 94.4 |
| Pergamino virus | Argentina | 82.4 | 74.4 | 79.3 | 93.9 |
| **Rio Mamore virus** | Perú/ Bolivia | 89.8 | 74.4 | 81.1 | 96.1 |
| Rio Mearim virus | Brazil | 85.4 | 74.4 | 79.9 | 96.1 |
| *S. aquaticus* LZ36 | Argentina | 82.5 | 66.7 | NA | NA |
| *S. aquaticus* LZ9 | Argentina | NA | NA | NA | NA |
| **Sin Nombre virus** | USA | 80.2 | 71.8 | 76.7 | 88.7 |
| Tunari virus | Bolivia | 86.2 | 71.8 | 79.2 | 95.2 |

S-segment fragment: 272 nts; M-segment fragment: 575 nts. Bold font: orthohantavirus named for the ICTV.

### Orthohantavirus infection in rodent populations

As the restrictions imposed due to the COVID-19 pandemic did not allow rodent capture campaigns, we analyzed rodent samples previously obtained for a study on the ecoepidemiology of leptospirosis conducted in the Center-Northern of the province of Santa Fe, which is a peripheral edge of the HPS endemic area of the Central region. We analyzed serum samples from 112 rodents of which 15 were synanthropic Murinae (12 *Mus musculus*, 2 *Rattus rattus* and 1 *R. norvegicus*) and 97 were native Sigmodontinae (21 *Akodon azarae, 53 Scapteromys aquaticus, 1 Holochilus chacarius*, 20 *Oligoryzomys flavescens*, 2 *O. nigripes*). From all rodents analyzed we only found reactivity in individuals of *S. aquaticus*, in which 18.9% had high IgG titers against hantavirus (Table 3). None of the individuals of the genus *Oligoryzomys* were seropositive. Seropositivity was significantly associated with the environmental setting, with all of the seropositive individuals trapped in one of the sites with higher human disturbance, and with the species *S. aquaticus* (Table 3). Among the infected individuals, 70% were females, 80% were sexually mature and 90% had either a low or moderate body condition (Table 4).

**Table 3.**
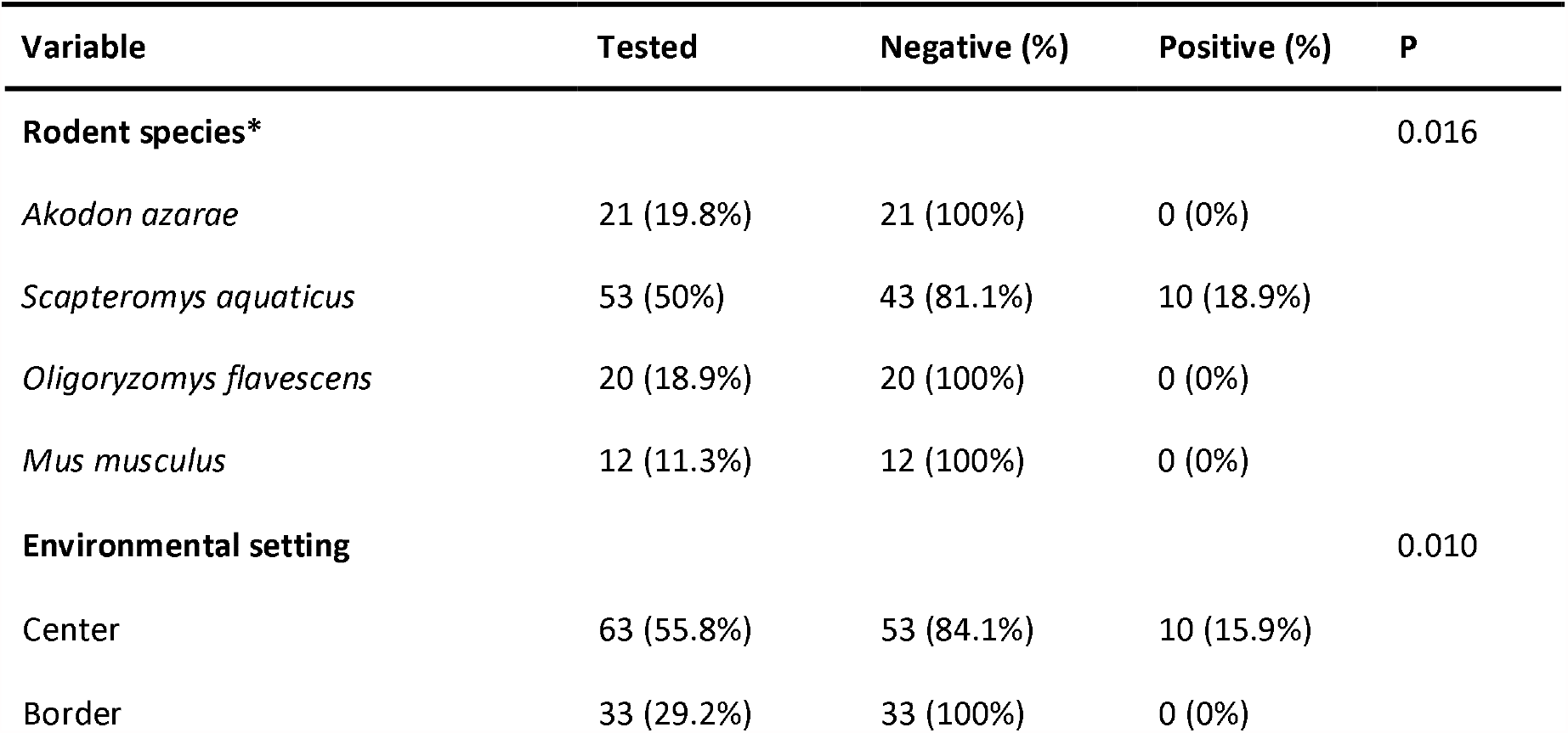
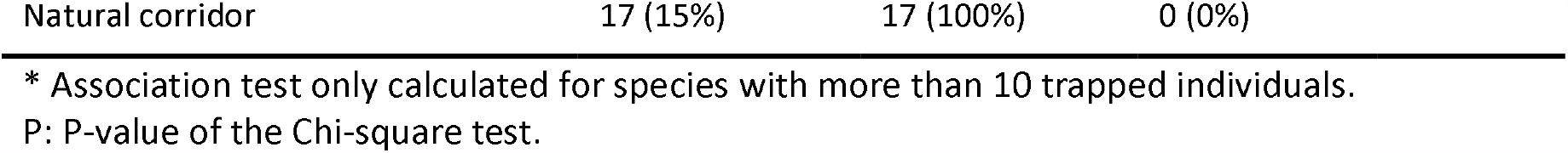
Hantavirus seroprevalence in murid and sigmodontine rodents from riverside settlements of the Paraná flooded savanna, Santa Fé, Central Argentina, from 2014 to 2015.

| Variable | Tested | Negative (%) | Positive (%) | P |
| --- | --- | --- | --- | --- |
| **Rodent species\*** |  |  |  | 0.016 |
| *Akodon azarae* | 21 (19.8%) | 21 (100%) | 0 (0%) |  |
| *Scapteromys aquaticus* | 53 (50%) | 43 (81.1%) | 10 (18.9%) |  |
| *Oligoryzomys flavescens* | 20 (18.9%) | 20 (100%) | 0 (0%) |  |
| *Mus musculus* | 12 (11.3%) | 12 (100%) | 0 (0%) |  |
| **Environmental setting** |  |  |  | 0.010 |
| Center | 63 (55.8%) | 53 (84.1%) | 10 (15.9%) |  |
| Border | 33 (29.2%) | 33 (100%) | 0 (0%) |  |

|  |  |  |  |
| --- | --- | --- | --- |
| Natural corridor | 17 (15%) | 17 (100%) | 0 (0%) |

\* Association test only calculated for species with more than 10 trapped individuals. P: P-value of the Chi-square test.

**Table 4.** RT-PCR result and IgG titers against Hantavirus in *Scapteromys aquaticus* from riverside settlements of the Paraná flooded savanna, Santa Fe, Central Argentina, from 2014 to 2015.

| ID | Sampling session | Sex | Reproductive status | Body condition | IgG titer | RT-PCR result |
| --- | --- | --- | --- | --- | --- | --- |
| LZ09 | Spring 2014 | Female | Mature | Low | 6400 | POS |
| LZ10 | Spring 2014 | Female | Immature | Moderate | 102400 | NEG |
| LZ11 | Spring 2014 | Female | Immature | Moderate | 51200 | NEG |
| LZ20 | Spring 2014 | Female | Mature | Moderate | 25600 | NEG |
| LZ34 | Autumn 2015 | Female | Mature | Low | 400 | NEG |
| LZ36 | Autumn 2015 | Male | Mature | High | 25600 | POS |
| LZ39 | Autumn 2015 | Female | Mature | Low | 1600 | NEG |
| LZ46 | Spring 2015 | Male | Mature | Moderate | 6400 | NEG |
| LZ63 | Spring 2015 | Male | Mature | Low | 6400 | NEG |
| LZ74 | Spring 2015 | Female | Mature | Low | 102400 | NEG |

Although tissue samples were not appropriately stored for the recovery of RNA, viral genome amplification was possible in two individuals, from which only short sequences could be recovered. A 488bp fragment corresponding to the S segment (nts: 430-918) and another of 444bp (nts: 8-452) from the M segment were obtained from rodent LZ09. An additional fragment of 283bp from the S segment (nts: 73 - 356) could be recovered from LZ36. Pairwise nucleotide and amino acid sequence analysis with all hantaviruses available in public repositories showed that this virus has not been characterized previously. Nucleotide identities ranged from 73.3% to 81.9% and from 66.2% to 73.2% for the S- and M- segment, respectively. Amino acid identities ranged from 86% to 93.5% and 68.9% to 76.3% for nucleoprotein (N) and Amino terminal Glycoprotein (Gn) respectively (S1 Table).

### Phylogenetic analysis

To determine the phylogenetic relationships among the novel hantaviruses described here with previously published orthohantaviruses (S2 Table), phylogenetic trees based on 26 sequences of the S- and M- segments were reconstructed using Bayesian inference. The virus amplified from the HPS case clustered together with APV, and both trees, based on the S- or M-segments, showed very similar topologies (Fig 2 and S1 Fig). Notably, this phylogroup occupied a separated cluster from the ANDV-like viruses previously described in Argentina. Regarding the viral sequences obtained from *S. aquaticus*, they showed striking differences regarding other orthohantaviruses from the Americas (Fig 2 and S2 Fig) and the trees corresponding to the S- and M-segments presented different topologies according to the fragments analyzed. We were only able to amplify fragments from the rodents LZ09 and LZ36 from different positions of the nucleoprotein-coding region, S-segment. The one from LZ36, which corresponded to the most conserved region of this segment, formed a monophyletic group together with Central and North American viruses (Choclo and Sin Nombre viruses), outside of all other South American clades (Fig 2). However, according to the M-segment, corresponding to the Gn-coding region, this novel virus seems to form a monophyletic group with the ANDV-like viruses (S2 Fig).

**Figure 2.**
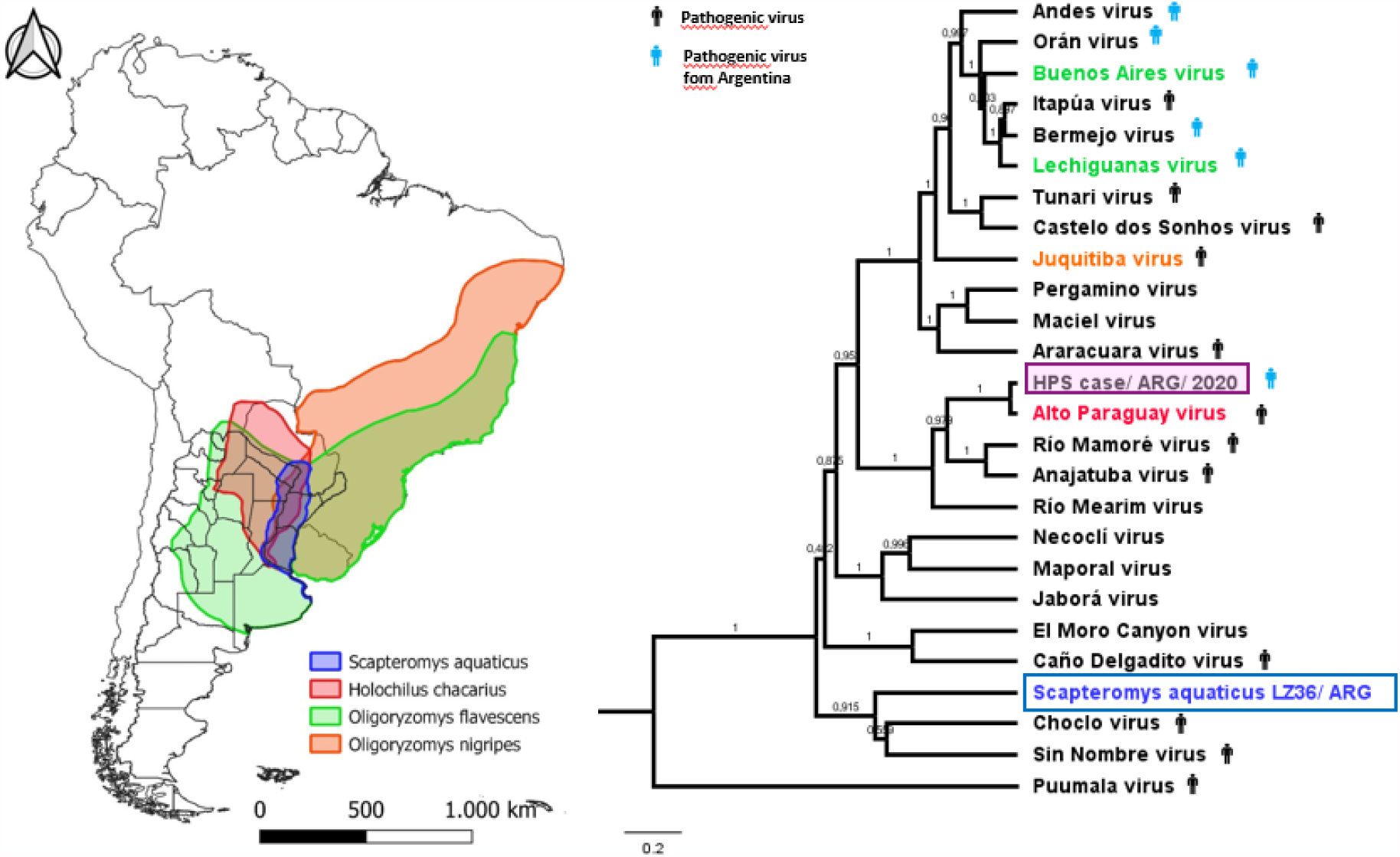
Geographical distribution of reservoir hosts present in the Hantavirus Pulmonary Syndrome endemic area of Central Argentina, and phylogenetic relationships between the viruses characterized here and selected orthohantaviruses from America. In the phylogenetic tree, the viruses present in the area are highlighted according to the color of its reservoir distribution shown in the map. Human figures in black indicate pathogenic viruses, human figures in blue indicate pathogenic viruses identified in Argentina. Map created using QGIS 3.14 Pi (QGIS Development Team); vector layers of the species distribution downloaded from the International Union for Conservation of Nature (IUCN). Phylogenetic relationships among South American Hantavirus genotypes based on Bayesian analysis of 26 taxa dataset of the S-segment partial sequences (283 bp for S. aquaticus and 1,286 bp for the remaining taxa), after 10 million generations. The tree is rooted relative to the position of Puumala virus. The bar represents 0.3 substitutions per nucleotide position. Constructed by BEAST (version 1.10.1) the algorithm MCMC (Metropolis-coupled Markov chain), Branches numbers indicate posterior probability values for the Bayesian inference.

## Discussion

Knowledge of the diversity and distribution of orthohantaviruses in the region has increased over the last 2 decades. The major agents causing HPS in Argentina are ANDV-like viruses, where 9 viruses have been characterized from patients and reservoir rodents [10]. In addition, LNV was also identified in Northwestern Argentina associated with few cases and rodents [22-24]. Among the known pathogenic orthohantaviruses, ANDV- and LNV-like viruses were also found in Chile, Bolivia, Uruguay and Brazil [24-27]. Other viruses not-associated with HPS were also described in neighboring countries, Jaborá virus, Rio Mearim virus and APV in *Akodon montensis, H. sciureus* and *H. chacarius*, respectively [28]. We report here 2 viruses that were never detected in Argentina until now: APV and a novel orthohantavirus found in *S. aquaticus* rodents.

APV was characterized by partial genetic analysis from a HPS case diagnosed in the context of the COVID-19 pandemic. Interestingly, APV was not associated with human disease to date. It was previously characterized from *H. chacarius* in Alto Paraguay department, Western Paraguay [28], more than 1200km northwards from the location of the patient. The HPS patient reported in the present work was exposed to rodents in a rural area of Tostado, Santa Fe province, and had no travel history mainly due to the strict mobility restrictions associated with the Covid-19 pandemic. Although *H. chacarius* is present in Central Argentina, the rodent reservoir hosting APV in the area remains to be confirmed. APV was previously identified from only one individual of *H. chacarius* [28] thus; the possibility that APV was hosted by other rodent species should not be excluded. Some kilometers downstream of the Salado River and Colastine River, in the northernmost edge of the known HPS endemic region of Central Argentina, we identified a novel virus in the sigmodontine *S. aquaticus*. Preliminary genetic characterization analysis could not identify the virus associated with *S. aquaticus* as any other known hantavirus. Based on these results, we strongly suggest that this virus is a new orthohantavirus and propose to name it as Leyes virus, because all the positive rodents were captured in the margins of the Leyes stream.

HPS is generally associated with severe clinical courses showing case-fatality rates ranging from 23.9% to 30.8% in Central Argentina [7,8], therefore,hantavirus infection is not suspected when facing mild clinical presentations. If APV infection is associated with a mild or severe disease should be carefully analyzed in the future, when a greater number of cases infected with this virus could be identified. However, a mild clinical presentation could be the reason for underdiagnosis of HPS in the Dry Chaco region, where only 2 HPS cases were previously identified since 1996.

The high hantavirus seroprevalence (18.5%) in *S. aquaticus* suggests that this rodent species is a new hantavirus host. The identification of this new host evidences underreporting of orthohantavirus circulation in the region, which reinforces the hypothesis that HPS has been under-diagnosed in the area. *S. aquaticus* can be found in semi-aquatic habitats, such as swamps, water courses, or flooded areas with low vegetation of the argentine ecoregions of Humid Chaco, Paraná flooded savanna, Espinal, Pampas and Iberá, as well as in western Uruguay, southern Paraguay and southern Brazil [29-31] (Fig 2). This species was also found to be a reservoir of pathogenic leptospires in areas of the Paraná flooded savanna from the provinces of Santa Fe and Buenos Aires [32, 14, 15]. The knowledge gaps on lack of evidence about the role of some rodent species as reservoirs hosts of these hantaviruses, favors the risk of orthohantavirus transmission and hinders HPS prevention. Environmental disturbances, such as floods and anthropogenic perturbations, may affect the geographic distribution, abundance and dynamics of rodent species as well as contact between humans and rodents [33, 34].

The main limitations of this work were the impossibility to get blood samples from the HPS patient and fresh samples from rodents. The limited amount of serum obtained from the patient and the tissue samples of rodents that were not appropriately stored for RNA recovery, only allowed us to obtain partial viral genetic sequences. However, our findings are conclusive and robust to confirm the viral identity of these 2 orthohantavirus circulating in Central Argentina.

Hence, in order to enhance prevention strategies, hantavirus surveillance must be taken as a public health priority focusing on the identification of new pathogenic viruses and on the monitoring of rodent hosts. Finally, it is important to determine if the detection of APV was due to the increased surveillance and social awareness due to the COVID-19 pandemic, or if this constitutes a relatively recent introduction of the pathogen from neighboring countries. Regarding Leyes virus, it would be important to determine if it is being implicated in human disease. According to the phylogenetic analysis based on partial sequences previously published from APV [28, 30] and those analyzed in the present study, both APV and Leyes virus could be considered as new orthohantavirus species. Further efforts are currently being undertaken to obtain full genomic sequences of these 2 new orthohantaviruses that could be used to update the diagnostic tools currently available.

In summary, despite HPS being described in Argentina 25 years ago and that the gained medical experience have helped to significantly reduce the case-fatality rates, HPS remains an emerging disease, as evidenced by the appearance of new viruses worldwide. Our findings implicate an epidemiological warning regarding these new orthohantaviruses circulating in Central Argentina as well as new rodent species that must be considered as hosts from now on.

## SUPPLEMENTARY TABLES AND FIGURES

**Table S1 – Pairwise nucleotide and amino acid identity percentages between virus sequences** amplified from the 2 *S. aquaticus*, Santa Fe, Argentina and selected American orthohantaviruses.

**Table S2 – GenBank Accession Numbers**.

**Figure S1. Phylogenetic relationships between the amplified sequences from a Hantavirus Pulmonary Syndrome case found in Argentina with selected American orthohantaviruses**. Trees were based on Bayesian analysis of partial M-segment sequences, after 10 million generations for a 696bp fragment of the Gn/Gc-coding region of Alto Paraguay virus/ ARG/ 2020 (nts: 2219 to 2915) and 25 complete sequences for the remaining taxa. The trees were rooted relative to the position of the Puumala virus. The bar represents 0.3 substitutions per nucleotide position. Constructed by BEAST (version 1.10.1) the algorithm MCMC (Metropolis-coupled Markov chain), Branches numbers indicate posterior probability values for the Bayesian inference.

**Figure S2. Phylogenetic relationships between the amplified sequences from a *Scapteromys aquaticus* rodent found in Argentina with selected American orthohantaviruses**. Trees were based on Bayesian analysis of partial M-segment sequences, after 10 million generations for a 444bp fragment of the Gn-coding region of *S. aquaticus* LZ9 (nts: 8 to 452) with 24 taxa dataset of partial sequences. The trees were rooted relative to the position of the Puumala virus. The bar represents 3 substitutions per nucleotide position. Constructed by BEAST (version 1.10.1) the algorithm MCMC (Metropolis-coupled Markov chain), Branches numbers indicate posterior probability values for the Bayesian inference.

