## Supplementary material for "Emerging Hantaviruses in Central Argentina: first case of Hantavirus Pulmonary Syndrome caused by Alto Paraguay Virus and a novel orthohantavirus in *Scapteromys aquaticus* rodent": Figure S2

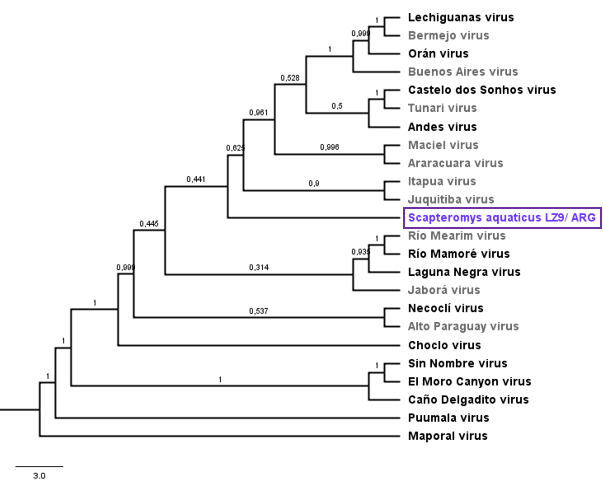

**Figure S2. Phylogenetic relationships between the amplified sequences from a *Scapteromys aquaticus*  rodent found in Argentina with selected American orthohantaviruses**. Trees were based on Bayesian analysis of partial M-segment sequences, after 10 million generations for a 444bp fragment of the Gn-coding region of *S. aquaticus* LZ9 (nts: 8 to 452) with 24 taxa dataset of partial sequences. The trees were rooted relative to the position of the Puumala virus. The bar represents 3 substitutions per nucleotide position. Constructed by BEAST (version 1.10.1) the algorithm MCMC (Metropolis-coupled Markov chain), Branches numbers indicate posterior probability values for the Bayesian inference.
