## Supplementary material for "Emerging Hantaviruses in Central Argentina: first case of Hantavirus Pulmonary Syndrome caused by Alto Paraguay Virus and a novel orthohantavirus in *Scapteromys aquaticus* rodent": Figure S1

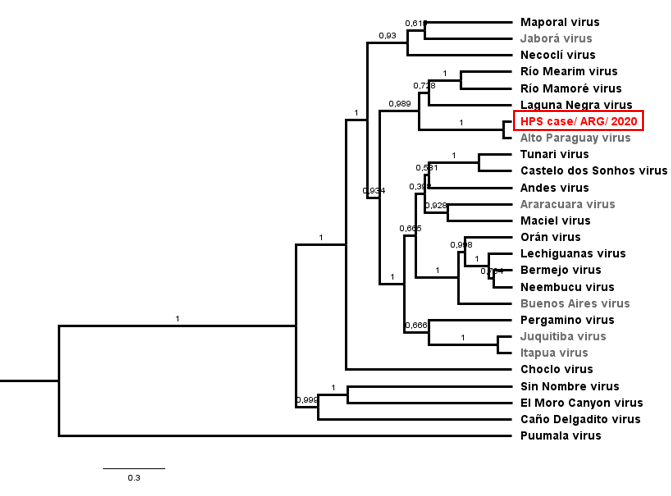

**Figure S1.** **Phylogenetic relationships between the amplified sequences from a Hantavirus Pulmonary Syndrome case found in Argentina with selected American orthohantaviruses**. Trees were based on Bayesian analysis of partial M-segment sequences, after 10 million generations for a 696bp fragment of the Gn/Gc-coding region of Alto Paraguay virus/ ARG/ 2020 (nts: 2219 to 2915) and 25 complete sequences for the remaining taxa. The trees were rooted relative to the position of the Puumala virus. The bar represents 0.3 substitutions per nucleotide position. Constructed by BEAST (version 1.10.1) the algorithm MCMC (Metropolis-coupled Markov chain), Branches numbers indicate posterior probability values for the Bayesian inference.
