## Supplementary material for "Emerging Hantaviruses in Central Argentina: first case of Hantavirus Pulmonary Syndrome caused by Alto Paraguay Virus and a novel orthohantavirus in *Scapteromys aquaticus* rodent": Table S2

**Table S2** – **GenBank Accession Numbers.**

|  | S-Segment | | M-Segment | |
| --- | --- | --- | --- | --- |
|  | Nucleotide | Amino acid | Nucleotide | Amino acid |
| Jaborá virus | GU205338 | ABC70874 | FJ409556 | ACR16367 |
| Sin Nombre virus | AF281850 | NP_941975 | KT885045 | ALI59819 |
| Pergamino virus | AF482717 | AAL82652 | AF028028 | AAB87914 |
| Maciel virus | AF482716 | AAL82651 | AF028027 | AAB87913 |
| Juquitiba virus | EU373729 | ABY76310 | AY963900 | AAX78361 |
| Bermejo virus | AF482713 | AAL82648 | AF028025 | AAB87911 |
| Lechiguanas virus | AF482714 | AAL82649 | AF028022 | AAB87908 |
| Orán virus | AF482715 | AAL82650 | AF028024 | AAB87910 |
| Buenos Aires virus | AF482711 | AAL82646 | AF028023 | AAB87909 |
| Itapua virus | EU373733 | ABY76314 | AY515601 | AAS00659 |
| Andes virus | AF324902 | NP_604471 | AF324901 | AAK14322 |
| Laguna Negra virus | AF005727 | YP_009506656 | NC_038506 | YP_009506658 |
| Alto Paraguay virus | DQ345762 | ABC70872 | Gn: AY515597 | Gn: AAS00655 |
| Alto Paraguay virus | DQ345762 | ABC70872 | Gc: AY515602 | Gc: [AAS00660](https://www.ncbi.nlm.nih.gov/protein/41351911) |
| Rio Mamoré virus | U52136 | AAC58450 | FJ608550 | ACU46022 |
| Rio Mearim virus | DQ451828 | ABE68627 | JX443701 | AFV36409 |
| Castelo Do Sonhos virus | AF307324 | AAG24912 | JX443702 | AFV36410 |
