## Supplementary material for "Emerging Hantaviruses in Central Argentina: first case of Hantavirus Pulmonary Syndrome caused by Alto Paraguay Virus and a novel orthohantavirus in *Scapteromys aquaticus* rodent": Table S1

**Table S1** – **Pairwise nucleotide and amino acid identity percentages between virus sequences amplified from the 2 *S. aquaticus*, Santa Fe, Argentina and selected American orthohantaviruses.**

| **Virus** | **Country** | **S-segment** | | | | **M-segment** | |
| --- | --- | --- | --- | --- | --- | --- | --- |
|  |  | **Nucleotide** | | **Amino acid** | | **Nucl.** | **Amin.** |
|  |  | **LZ9** | **LZ36** | **LZ9** | **LZ36** | **LZ9** | **LZ9** |
| **Andes virus** | Argentina | 74.8 | 83.2 | 88.4 | 88.9 | 73.2 | 71.1 |
| Bermejo virus | Argentina | 81.8 | 83.9 | 87.6 | 87.5 | NA | NA |
| Buenos Aires virus | Argentina | 75.8 | 82.2 | 87.6 | 87.5 | 71.3 | 71.9 |
| **Lechiguanas virus** | Argentina | 77.5 | 84.6 | 87.6 | 87.5 | 71.9 | 72.6 |
| Maciel virus | Argentina | 74.2 | 82.8 | 86 | 87.5 | NA | NA |
| **Oran virus** | Argentina | 75.6 | 82.9 | 87.6 | 87.5 | 69.5 | 68.9 |
| Pergamino virus | Argentina | 74.6 | 81.5 | 88.4 | 87.5 | NA | NA |
| *S. aquaticus* LZ36 | Argentina | NA | 100 | NA | 100 | NA | NA |
| *S. aquaticus* LZ9 | Argentina | 100 | NA | 100 | NA | 100 | 100 |
| Juquitiba virus | Arg./ Brazil | 75.9 | 80.2 | 86.8 | 87.5 | NA | NA |
| **Laguna Negra virus** | Arg./ Parag./ Brazil | 76 | 82 | 86 | 87.5 | 71.7 | 68.9 |
| Tunari virus | Bolivia | 76.5 | 82.2 | 87.6 | 86.1 | NA | NA |
| Anajatuba virus | Brasil | 78.5 | 82.8 | 90.1 | 87.5 | NA | NA |
| Araracuara virus | Brasil | 77.7 | 81.1 | 88.4 | 87.5 | NA | NA |
| Jabora virus | Brazil | 76.9 | 75.8 | 90.9 | 91.6 | 72.7 | 76.3 |
| **Castelo Do Sonhos virus** | Brazil | 79.9 | 84.3 | 92.5 | 87.5 | 71.5 | 71.1 |
| Rio Mearim virus | Brazil | 77 | 83.9 | 88.4 | 88.9 | 72.6 | 68.9 |
| **Necoclí virus** | Colombia | 79.3 | 80 | 87.6 | 79.5 | 70.2 | 70.4 |
| **Choclo virus** | Panamá | 78.6 | 82.5 | 88.4 | 80.7 | 71.8 | 68.9 |
| Alto Paraguay virus | Paraguay | 76.8 | 82.1 | 90.9 | 86.1 | NA | NA |
| IP16 virus | Paraguay | 77.8 | 75.1 | 90.9 | 91.6 | NA | NA |
| Itapúa virus | Paraguay | 78.1 | 84.9 | 86.8 | 87.5 | NA | NA |
| **Rio Mamore virus** | Perú/ Bolivia | 78 | 82.5 | 90.1 | 87.5 | 71.1 | 72.2 |
| **El Moro Canyon virus** | United States | 80.1 | 79 | 81 | 91.7 | 67.8 | 56.6 |
| **Sin Nombre virus** | United States | 73.3 | 82.2 | 86 | 81.5 | 66.2 | 54.1 |
| **Caño Delgadito virus** | Venezuela | 77.2 | 78.8 | 89.5 | 77.8 | 68.3 | 55.2 |

S-segment: 488bp (430-918); M-segment: 444bp (nts: 8-452), from rodent LZ9. M-segment: 283bp (nts: 73 - 356) from LZ36.
